# WebSeq: A Genomic Data Analytics Platform for Monogenic Disease Discovery

**DOI:** 10.1101/2021.11.16.460500

**Authors:** Milind Agarwal, Kshitiz Ghimire, Joy D. Cogan, Undiagnosed Disease Network, Janet Markle

## Abstract

Whole exome sequencing (WES) is commonly used to study monogenic diseases. The application of this sequencing technology has gained in popularity amongst clinicians and researchers as WES pricing has declined. The accumulation of WES data creates a need for a robust, flexible, scalable and easy-to-use analytics platform to allow researchers to gain biological insight from this genomic data. We present WebSeq, a self-contained server and web interface to facilitate intuitive analysis of WES data. WebSeq provides access to sophisticated tools and pipelines through a user-friendly and modern web interface. WebSeq has modules that support i) FASTQ to VCF conversion, ii) VCF to ANNOVAR^1^ CSV conversion, iii) family-based analyses for Mendelian disease gene discovery, iv) cohort-wide gene enrichment analyses, (v) an automated IGV^2^ browser, and (vi) a ‘virtual gene panel’ analysis module. WebSeq Pro, our expanded pipeline, also supports SNP genotype analyses such as ancestry inference and kinship testing. WebSeq Lite, our minimal pipeline, supports family-based analyses, cohort-wide gene enrichment analyses, and a virtual gene panel along with the IGV^2^ browser module. We anticipate that the rigorous use of our web application will allow researchers to expedite discoveries from human genomic data^3^. WebSeq Lite, WebSeq, and WebSeq Pro are fully containerized using Docker^4^, run on all major operating systems, and are freely available for personal, academic, and non-profit use at http://bitly.ws/g6cn

## Introduction

An abundance of computational tools is available to researchers for genomics data analysis. However, availability of a resource may not translate into accessibility. Most genomic toolkits are built by experienced software developers, bioinformaticians, and computer scientists. Although these toolkits are powerful, most have subtle setup instructions and system dependencies that may be barriers to use for biologists who lack computer science training. In WebSeq, we have brought together several existing bioinformatics toolkits in one place, created new analysis capabilities, and provided an intuitive user interface.

A plethora of bioinformatics toolkits are available to facilitate analysis of genomic data. Some of these tools allow for robust SNP data analysis (ex. PLINK^5,6^, GATK^7^), while others allow for powerful variant annotation (ANNOVAR^1^). Most of these tools need to be run through command line, while some of the more recent toolkits have dedicated web servers (ex. wANNOVAR^3^, GeneMANIA^8^) to allow for easier analysis. For example, KING^9^ is a widely used toolkit primarily for relationship inference and flagging pedigree errors. KING^9^ is also implemented on command line, and lacks an accessible graphical user interface. However, KING’s^9^ fast and robust performance is very useful for a wide variety of research problems. PLINK^5,6^ is another popular toolkit used for genomic data analyses. It is open source and written in C/C++ which contributes to its scalability and computational efficiency. While offering a huge suite of services, PLINK^5,6^ runs completely through command line and this could discourage researchers without command line experience from taking advantage of this tool. Another example is popular genomic visualization-centric toolkit IGV^2^ which allows for interactive exploration of large, integrated genomic datasets. It supports a variety of data types, including array-based and next-generation sequence data, and genomic annotations. While IGV^2^ is high-performance and offers rich visualizations of the genome, manual variant inspection remains tedious.

WebSeq attempts to address some shortcomings of such toolkits and combine their strengths with our own analytics capabilities all together into one platform. WebSeq’s web-based interface can securely convert FASTQ file data into BAM and VCF formats using GATK^7^, SAMtools^10^, and BWA^11^. It can also annotate VCF file data through ANNOVAR^1^ and conduct family based analysis for Mendelian disorders. It is especially practical for identifying putative disease-causing variants in a family in which multiple family members have been sequenced. It can perform cohort level analyses such as finding genetic homogeneity, querying particular genes of interest, and conducting pathway enrichment analyses with GeneMANIA^8^. A built-in IGV^2^ Report generator allows the researcher automate the process of browsing through variants and minimize active time spent confirming variant calls. WebSeq also supports SNP genotype-based analyses, such as kinship inference through KING^9^, sex inference through PLINK^5,6^, and Principal Component Analysis (PCA) based ancestry inference. Refer to Figure 1 below for an overview of WebSeq’s modules, supported data formats, nested services, and memory requirements.

**Figure 1:**
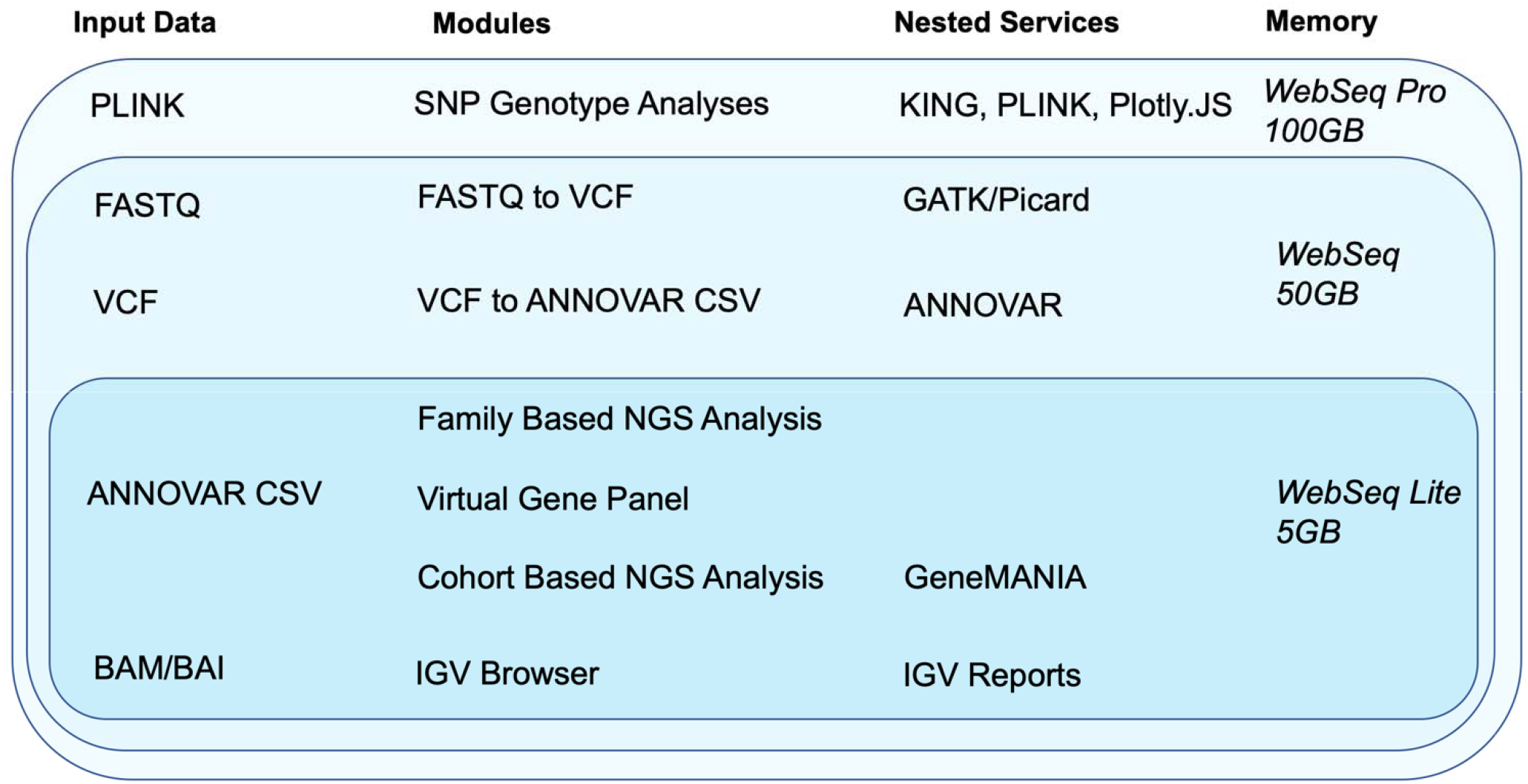
Modules, data input requirements, nested services and memory requirements for WebSeq Lite, WebSeq and WebSeq Pro.

## Materials and Methods

### Application Architecture

#### Backend Server

The backend is made entirely out of Flask^12^, a lightweight micro framework. Each functionality is offered as a separate view within well-defined modules. KING^9^, PLINK^5,6^, GATK^7^, and IGV^2^ Reports are nested within the backend architecture and are exposed to the user through the web interface.

#### Message Broker

WebSeq uses the Celery^13^ message broker along with the Redis database as a queueing system. This is essential for monitoring progress of long running tasks and ensuring that the application doesn’t become unresponsive. This also ensures that data intensive tasks can be run in the background in parallel with the web interface itself. In this manner, the message broker facilitates communication between the server and the client, as shown in Figure 2.

**Figure 2:**
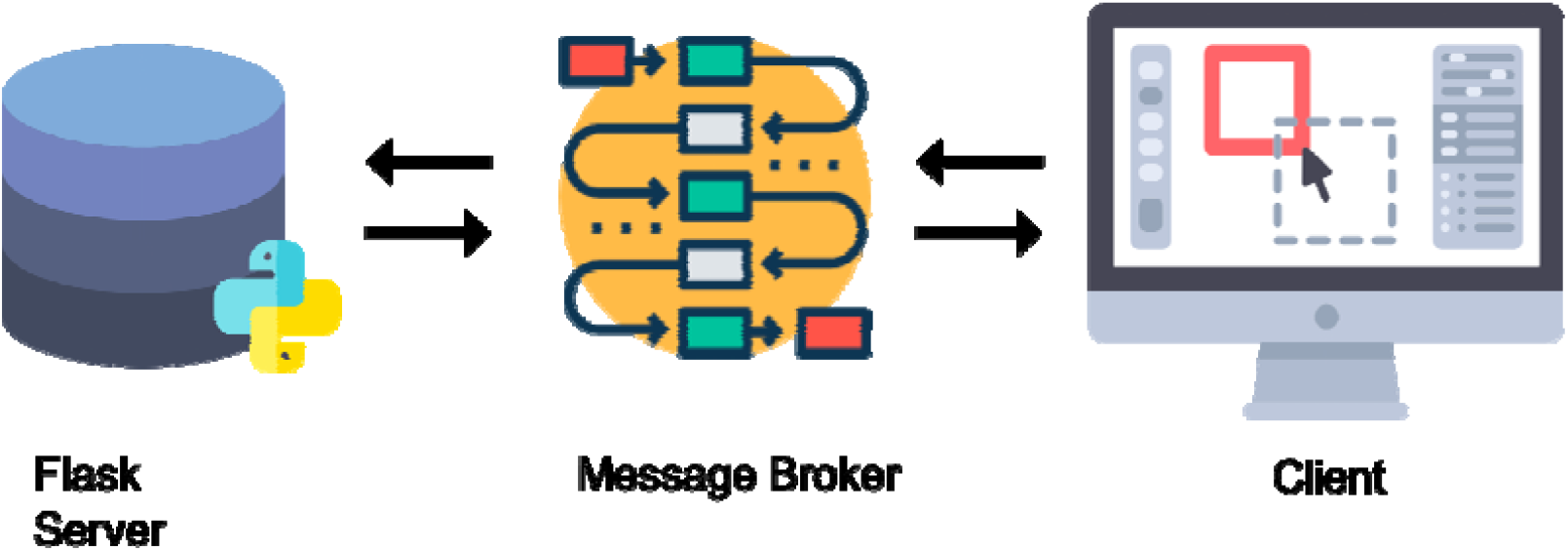
WebSeq Architecture. The backend consists of the Flask^12^ server and the nested services. The client communicates with the server through HTTP requests and a message broker.

#### Client Side/Frontend

The interface is built entirely out of the fundamental scripting and markup languages. We use HTML for structure of the web application, Bootstrap and custom CSS for styling, and JavaScript for interactivity and data streaming through HTTP requests. Any file you “upload” never leaves your computer. They are simply streamed to a temporary location inside the application sandbox for processing, and then streamed back to you for download via the web interface. We use custom AJAX requests to allow for streamlined communication, parsing of data and parameters, and download of result files through responses.

#### Docker^4^

The entire application is containerized using Docker^4^. Docker^4^ is a powerful toolkit that helps us package WebSeq with all the parts it needs, such as libraries and software dependencies, to run smoothly on any operating system or platform. It ensures that users do not need to perform complicated setup instructions to launch WebSeq.

#### Nested Services

ANNOVAR^1^: ANNOVAR^1^, and its web version wANNOVAR^3^, are popular choices for variant level annotation. ANNOVAR^1^ is a command line interface (CLI) tool written entirely in Perl. ANNOVAR allows researchers to annotate their VCF files with information from a wealth of curated databases. This facilitates further downstream data analysis since it allows users to make informed decisions about variants. In our VCF to ANNOVAR^1^ converter module, we utilize ANNOVAR^1^ scripts and expose them through a very intuitive easy to use front-end interface. This allows for robust, secure, in-house annotation without uploading VCF data to third-party servers.

GeneMANIA Pathway Visualizer^8^: GeneMANIA^8^ is a recent web-based analytics toolkit that allows the user to visualize relationships between a query of genes. We offer automatic linking to GeneMANIA^8^ on upload of a file with query loci. These can be visualized from within the web interface. This is extremely valuable in detecting potential pathways and getting information from related publications.

Integrated Genome Viewer: IGV^2^ is primarily a Java based Desktop application, but its creators have also released IGV^2^ Web, and a JavaScript plugin for web developers to incorporate into their projects. We incorporate IGV’s JavaScript plugin and IGV^2^ Reports into our application.

## Results

WebSeq offers 7 modules for variant annotation, and variant filtering in a self-contained web application, as shown in Figure 3. These modules can be used in two main workflows, monogenic disease discovery and SNP genotype analyses, both of which are outlined in Figure 4.

**Figure 3:**
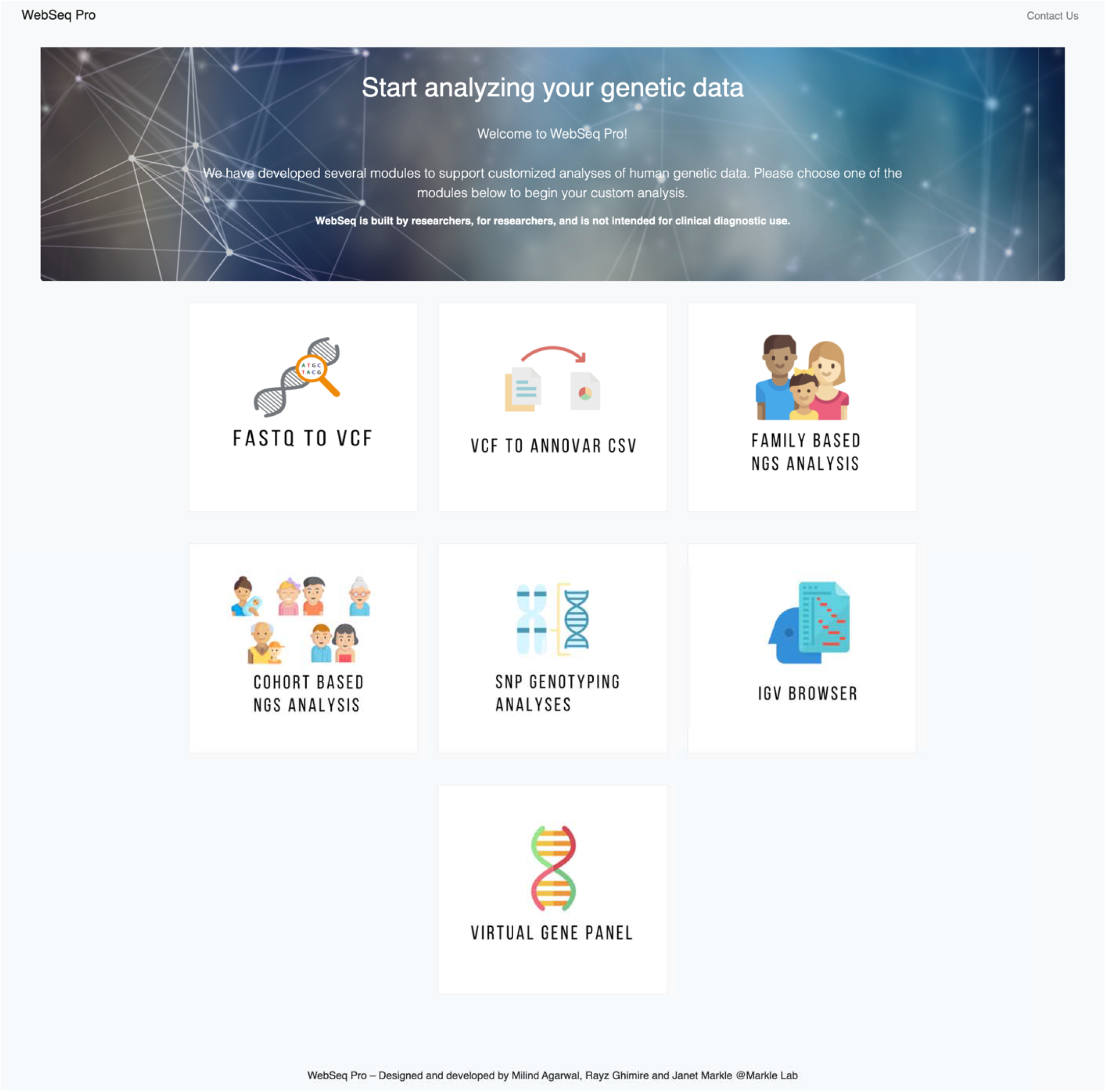
WebSeq Pro’s Home Page displays all 7 available modules: FASTQ to VCF, VCF to ANNOVAR^1^ CSV, Family Based NGS Analysis, Cohort Based NGS Analysis, SNP Genotyping Analyses, IGV^2^ Browser, Virtual Gene Panel.

**Figure 4:**
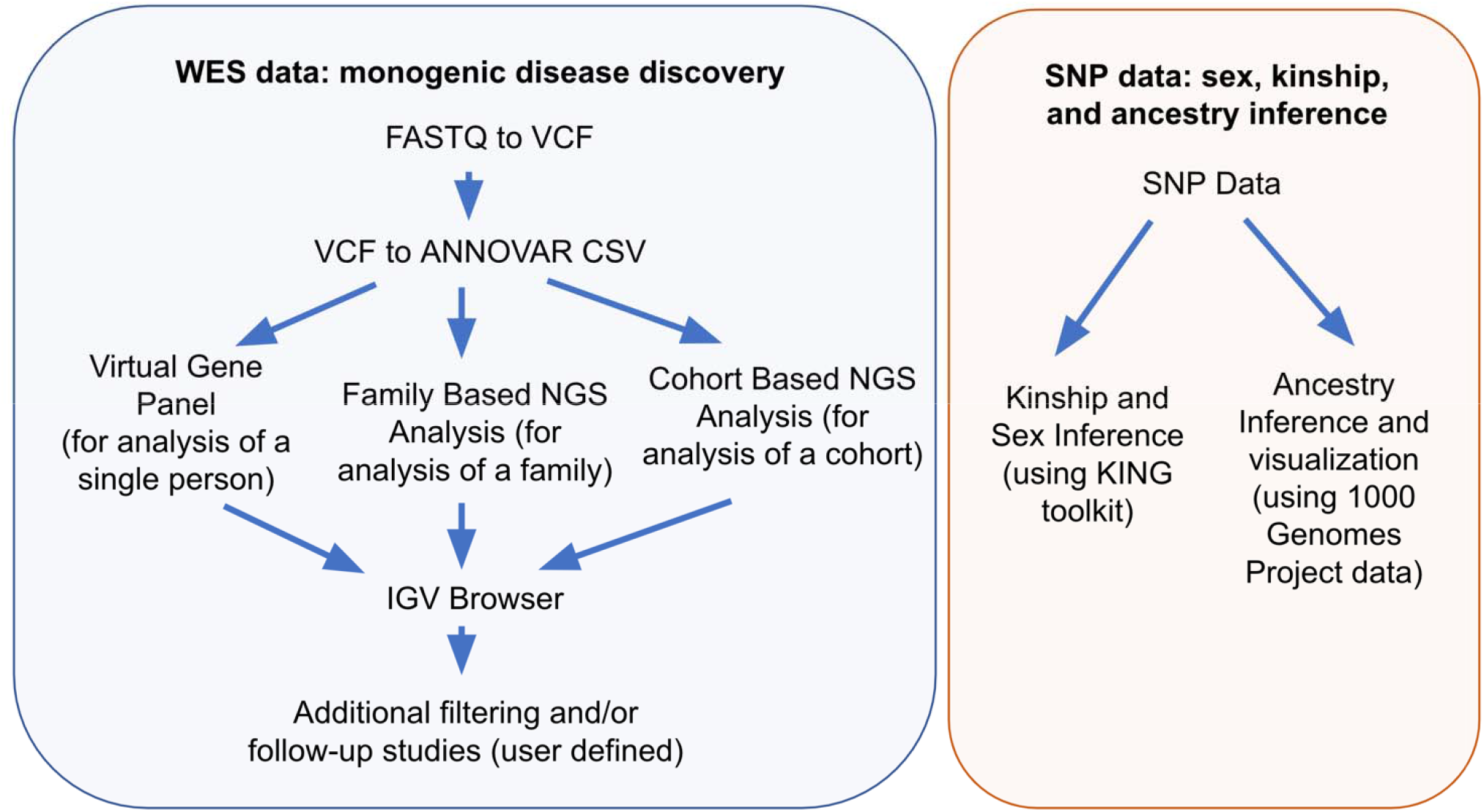
WebSeq workflows for monogenic disease discovery and SNP genotype analyses.

### Module 1) FASTQ to VCF

Starting with .fastq files from WES, this module will align the data to the GRCh38/hg38 reference genome, generate .sam and .bam files, mark duplicate reads, recalibrate base quality scores, and call variants. It will generate a VCF file for use in the next module.

1. For alignment, this module uses the BWA-MEM^11^ alignment algorithm for aligning sequence reads or long query sequences against a large reference genome, in this case the human GRCh38/hg38 reference genome.. The algorithm is robust to sequencing errors and applicable to a wide range of sequence lengths from 70bp to a few megabases.
2. Following deduplication, we use GATK^7^/Picard^14^ tools to sort the SAM file and then perform base quality score recalibration (BQSR^15^). This step corrects base quality scores in the data from systematic technical errors based on a set of known true variants, such as those generated from humans in the 1000 Genomes^16^ project.
3. We use GATK’s^7^ HaplotypeCaller program to call variants. For each sample, the HaplotypeCaller program uses the GRCh38/hg38 reference genome and BAM files to produce an output VCF file to be used in the next module.

### Module 2) VCF to ANNOVAR^1^ CSV

Annotating VCF files is a crucial first step before filtering and prioritizing variants. ANNOVAR^1^ is a toolkit that can functionally annotate genetic variant data with update-to-date information from databases. These annotations can be performed through wANNOVAR^3^ online or through the ANNOVAR^1^ command line. Our module abstracts away the need to use command line and also takes away the file size restriction of wANNOVAR^3^. This module allows for secure, and ANNOVAR^1^ server downtime-insensitive, annotation of VCF files. This module generates CSV files that can be used for further analysis in the other WebSeq modules.

### Module 3) Family Based NGS Analysis

#### 2a) Cleaning and Prefiltering

Genomic data demands high-quality metrics for subsequent filtering and analysis. However, too many annotations can increase your file sizes dramatically. Cleaning will reduce file sizes by removing columns from ANNOVAR^1^ files that are not typically used for downstream filtering and analysis, while preserving high quality features such as allele frequencies from gnomAD^17^, and results from variant effect predictors like SIFT^18^, Polyphen-2^19^ and CADD^20^.

Reducing your search domain may be useful for genomic data analyses including WES analysis. For instance, in a particular study, you might want to look at only exonic and splicing variants. The prefiltering module allows you to remove certain variants based on whether they fall in coding regions, intergenic regions, 5’ or 3’ untranslated regions, etc. You can use this step to select the categories of variants you’d like to keep, and the application filters out all other variant types in your data.

#### 2b) Annotation

Additional gene-level annotations can be added, to aid in subsequent filtering of variants according to gene-level tolerance to mutation. Currently, this module will add the following annotations to your data:

1. Gene Damage Index (GDI)^21^: The gene damage index (GDI)^21^ is the accumulated mutational damage of each human gene in healthy human population, based on the 1000 Genomes Project^16^ database (Phase 3) and of the CADD^20^ score for calculating impact. Since highly damaged human genes are unlikely to be causal for severe monogenic diseases, GDI^21^ can be used to filter out variants present in highly damaged (high GDI^21^) genes^22^.
2. Mutation Significance Cutoff (MSC)^23^ The MSC^23^ score of a gene represents the lowest expected clinically/biologically relevant CADD^20^ cutoff value for that specific gene. For each gene, a variant with high phenotypic impact (i.e. possibly damaging) is any variant with a CADD^20^ score equal or above the MSC^23^, and low phenotypic impact (i.e. benign) is any variant with a CADD^20^ score below the MSC^23,24^.
3. pLI, pRec, pNull scores^25^ The pLI^25^ score is the probability that a given gene falls into the Haploinsufficient category, therefore is extremely intolerant of loss-of-function variation. Genes with high pLI^25^ scores (pLI^25^ ≥ 0.9) are extremely LoF intolerant, whereas genes with low pLI^25^ scores (pLI^25^ ≤ 0.1) are LoF tolerant.
4. Closest Disease Causing Gene (CDG^26^) To facilitate the discovery of novel gene-disease associations, the CDG^26^ database and server provide the putative biologically closest known disease-causing genes (and their associated phenotypes) for 13,005 human genes not reported to be disease-causing.

#### 2c) Filtering by Allele Frequency

Next, you might want to filter your data by allele frequency. User may specify an allele frequency cutoff based on data from either WES or whole genome sequencing data in the gnomAD^17^ database. For instance, setting a cutoff of gnomAD^17^ MAF < 0.01 would allow you to restrict your search to only rare genetic variants with minor allele frequency 0.01 (or 1%) or less in the gnomAD^17^ database. This step is very useful to further narrow your search domain.

#### 2d) Modes of Inheritance

This module is useful for narrowing your search to variants that follow a specified genetic mode of inheritance within a given family, provided you have WES data from multiple family members. Here, the user can choose from a variety of zygosities (homozygous, heterozygous or absent) for each individual to test various inheritance models. Since you’ll most likely want to inspect the results of this file manually, you can also annotate the results of this step with HGNC^27^ Gene Names. The application will automatically find the full HGNC^27^ gene names, and report synonyms and aliases if the official gene name is not found.

Prior to conducting this step, it is highly recommended that you clean, prefilter, annotate, and filter (by minor allele frequency) your data. Since this step essentially does pairwise comparisons for each variant, preprocessing your data according to the workflow will greatly speed up computation.

Example: Let’s say you have processed data from three individuals: PATIENT (male), MOTHER, and FATHER for a trio-based study. First, upload the filtered .csv files containing vairants from each person in this trio. Next, specify the genetic mode of inheritance you wish to test. Table 1 below specifies the parameters you could choose in order to narrow your search to variants following various genetic modes of inheritance. The user may only test one mode of inheritance at a time. However, you may re-load the Modes of Inheritance module, re-upload the filtered .csv files, and then iteratively test different modes of inheritance.

**Table 1:** Examples of possible search parameters for a trio, that can be tested with the ‘Modes of Inheritance’ module.

| PATIENT | MOTHER | FATHER | X-Rec | CompHet | Hypothesis Tested |
| --- | --- | --- | --- | --- | --- |
| Het | Absent | Absent | - | - | *de novo* mutation |
| Het | Absent | Het | - | - | Autosomal Dominant (inherited from father) |
| Hom | Het | Het | - | - | Autosomal Recessive, homozygous mutation |
| Het | Het | Het | - | YES | Autosomal Recessive, compound heterozygous mutation |
| Hom | Het | Absent | YES | - | X-linked Recessive<br>Note: Hemizygous variants are annotated as homozygous through ANNOVAR<sup>1</sup> |

### Module 4) Cohort Based NGS Analysis

#### 3a) Unbiased Cohort Analysis

For cases when researchers have incomplete family medical histories, and therefore don’t have a high level of certainty about the mode of inheritance in a given family, this module can be extremely valuable. This step will analyze your cohort data to find genetic homogeneity. It produces a ranked file with the most recurring genes on top, i.e. genes that have variants in the most people will show up on top. Since these files can become huge in size and can overload memory, all computation for this module is parallelized on the server. It is strongly recommended that only files that have undergone some preliminary filtering (e.g. filtered by minor allele frequency) be used an input for this module.

#### 3b) Biased Cohort Analysis

This tool is a query-based form of cohort analysis. This module collects all instances of mutations in a particular query gene across your entire cohort. For example, if you had a cohort of 100 individuals with inflammatory bowel disease, and you wanted to find out how many individuals had a mutation in the gene *NOD2*, this module would be the right tool to use. Through a friendly user interface, this module will produce a file NOD2.csv with all mutations present in this gene in your entire cohort, along with identifiers mapping back to the person each *NOD2* mutation was found in.

#### 3c) Variant Prioritization with Blacklists

This module is inspired by the idea for blacklisting cohort-specific variants in ReFiNE (Reducing Fal e-positives in NGS Elucidation)^28^. We extend its ideas by allowing for upload of ANNOVAR^1^ formatted files. In this module, you can generate your own in-house blacklist of variants, and then also apply this private blacklist to any file from the workflow. The module will exclude any variants that appear in the blacklist from your data. Since cohort sizes for generation of blacklists might be different between research labs, we allow the user to choose a tolerance before generating their own blacklist. Tolerance is defined as the minimum percentage of people in your cohort that a variant must be present in, in order to qualify as a blacklisted variant.

#### 3d) GeneMANIA^8^ Pathway Analysis

This module allows you to upload a file from within the WebSeq workflow, and visualize the network formed by unique genes present in your file in GeneMANIA^8^. GeneMANIA^8^ is very easy to use and is quick to produce high-quality networks for queries. You can use GeneMANIA^8^ to find new members of a pathway or complex, find additional genes you may have missed in your screen or find new genes with a specific function, such as protein kinases. Your question is defined by the set of genes you input^29^. See Figure 5 for an example cohort-based analysis workflow that includes unbiased cohort analysis, variant prioritization and pathway analysis.

**Figure 5:**
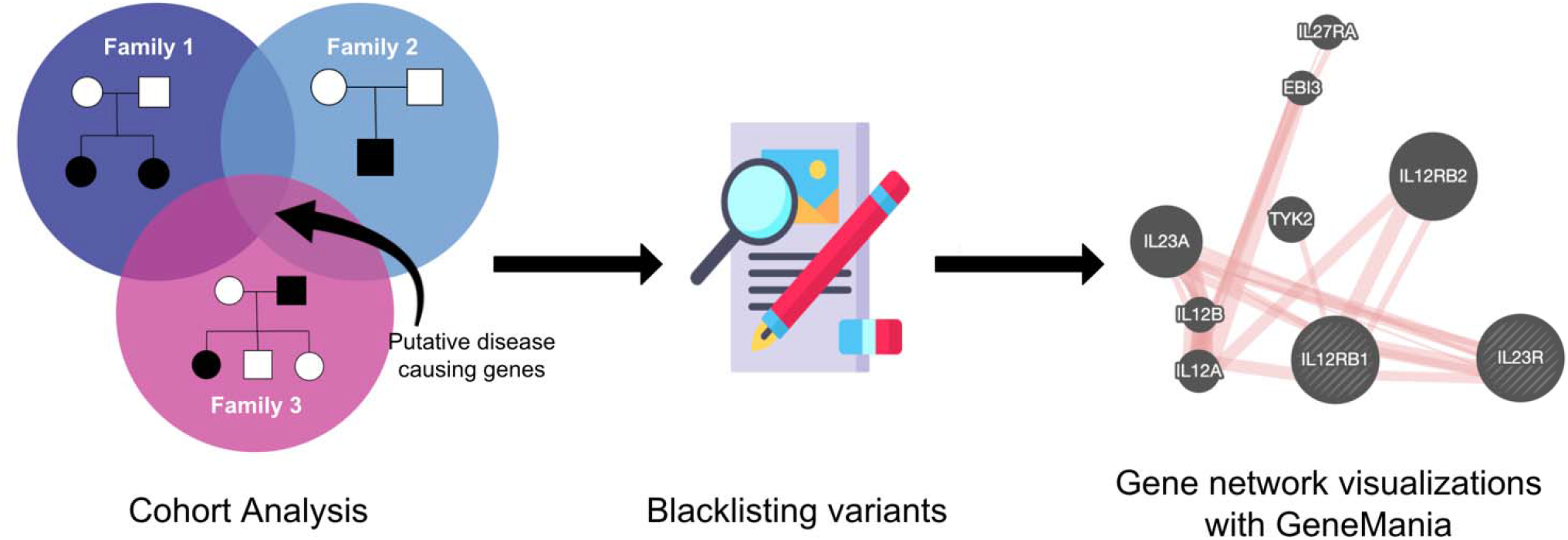
Cohort Based NGS Analysis Example Workflow. Unbiased cohort analysis to find putative disease-causing variants, followed by removal of in-house blacklisted variants, and visualization of gene network with GeneMANIA^8^.

### Module 5) Virtual Gene Panel

In some instances, researchers may have *a priori* knowledge about the genes that underly a certain phenotype. This module allows you to easily extract variants in genes of interest (from a user-defined list) from an individual’s CSV file. To use this module, the user must first generate a CSV file with the column heading Gene.refGene and with all genes of interest listed in that column. Make sure to use gene names, as per the GRCh38/hg38 reference genome, which may differ from the corresponding protein names. Using this file, the Virtual Gene Panel module can be used to generate a list of variants present in an individual’s WES data within these genes of interest only. If the user wishes to focus only on rare variants in genes of interest and is analyzing data from only a single person (e.g. proband), we recommend using this module after applying a Minor Allele Frequency filter. This module is flexible and may be applied to other results files generated by WebSeq, such as the results of a Family Based Analysis, or a Cohort Analysis.

### Module 6) SNP Genotyping Analyses

#### 4a) Kinship and Sex Inference

Through this module, we expose a graphical user interface to the robust and extremely fast command line tool KING^9^. KING^9^ can be used to check family relationship and flag pedigree errors by estimating kinship coefficients and inferring Identity by Descent (IBD) segments for all pairwise relationships^30^. KING^9^ provides a kinship coefficient for each pairwise comparison, and this score can be used to interpret the degree of relatedness between individuals. KING^9^ is extremely fast (seconds to identify all close relatives in 10,000s of samples)^30^. This can be an extremely valuable tool for exploratory analyses, quality control, and identification of unknown relationships between individuals within your cohorts.

The PLINK^5,6^ toolkit offers a wealth of different functionalities such as data management, basic statistics, linkage disequilibrium calculation, IBD matrix calculations for relationship inference, population stratification using PCA, association analyses for genome-wide association studies etc^5^. WebSeq’s sex inference module uses the PLINK^5,6^ toolkit to find the genetically inferred sex of each individual in your dataset. If there’s a conflict, the individual is marked as such in the output. By default, our interactive user interface abstracts away the need to compose Bash commands or use command line to use the PLINK^5,6^ toolkit and allows you to select the mode of sex inference: X-chromosome, and Y-chromosome data.

#### 4b) Ancestry Inference

Accurate inference of genetic ancestry is of fundamental interest to many biomedical, forensic, and anthropological research areas. Genetic ancestry may relate to genetic disease risks, and self-reported ancestries in studies can be inaccurate. In a genome wide association study, failing to account for differences in genetic ancestry between cases and controls may also lead to false-positive results^31^.Genetic markers allow for accurate and reliable inference of an individual’s ancestral origins. In this toolkit, we allow the user to visualize ancestries of their sample data with the 1000G^16^ data as reference via PCA. We offer pre-merged datasets for all chromosomes to speed up processing. The principal components are calculated behind the scenes through PLINK^5,6^ and are visualized through an embedded interactive scatter plot through PlotlyJS^32^.

### Module 7) Integrated Genome Viewer (IGV)^2^ Browser

Once you have a list of variants that pass your desired filtering criteria (e.g. after analysis in the mode of inheritance module, virtual gene panel module, or cohort analysis module), you may wish to visually inspect the aligned sequencing reads for quality and reliability. IGV^2^ requires you to enter a query locus, and then manually make note of whether to choose the variant as a potential candidate for downstream analyses. This is usually a time consuming, error-prone, and impractical step for high throughput data applications. We leverage the command line toolkit IGV^2^ Reports which generates a standalone report of all the variants in a particular sequence of search queries. This requires use of command line and preparation of BED/VCF formatted search queries. We allow users to utilize this tool from within the web interface, by selecting the appropriate BAM files, and the query file (CSV) from a previous step. The user can ‘browse’ through these variants by clicking YES/NO/MAYBE buttons. This alleviates the need to copy and paste one locus at a time. Once the user has gone through all the variants, the original file along with the responses annotated is made available for download.

### Validating WebSeq Functionality: Data from the Undiagnosed Disease Network

To test the performance of WebSeq, we obtained data from 6 patients and their parents (i.e., 6 ‘trios’) previously analyzed by WES by the Undiagnosed Disease Network (UDN)^33^, Vanderbilt University site. UDN investigators had used trio-based genome-wide sequencing analysis employing commercially available software tools to reach a monogenic disease diagnosis for each patient. In 5 of the 6 patients, a single nucleotide variant or small insertion/deletion variant, both detectable by WES, had been determined by the UDN to be disease-causing. The sixth patient was known to have a 1.66kb deletion, not detectable by standard WES data processing methods. A complete re-analysis of each of these 6 trios was done using WebSeq. First, .fastq files were converted to VCF, and .bam and .bai files were generated. Next, variant annotation was performed, then variants were pre-filtered to retain only those affecting coding or splicing sites. Variants were further filtered using MAF threshold values provided in Table 2. The 3 resulting files for each trio were then analyzed according to the genetic mode of inheritance shown in Table 2. In the 2 patients known to have *de novo* disease-causing mutations, variants underwent further manual filtering to retain only private alleles, i.e. alleles absent from gnomAD reference datasets. Next, variant calls were verified by inspecting the raw sequencing reads using .bam and .bai files and the WebSeq IGV browser module. Finally, the candidate variants generated by this WebSeq analysis were compared against the known causal mutations identified by the UDN. The 1.66kb deletion present in the patient from Trio 6 was not readily identified using this analysis pipeline, as expected. Alternative genome-wide sequencing approaches and/or data analysis pipelines may be needed for the detection of large copy number variants, non-coding variants, repeat expansions, or DNA methylation disorders^34^. For the 5 trios with SNV or small insertion/deletion variants, the causal mutation was identified by WebSeq analysis in all cases (Table 2). The rich variant annotation provided by the WebSeq pipeline also enables additional variant filtering by user-defined preferences, such as the prioritization of variants with high CADD scores. When each patient’s candidate variants were ranked using this approach, the causal mutation was the top candidate in all patients from Trios 1, 2, 3, and 5, and in Trio 4, the causal mutation was 1 of only 3 identified frameshift mutations (Table 2). These results demonstrate the utility of WebSeq for efficient analysis of trio-design WES data, and indicate that WebSeq can perform comparably to commercially available data analysis pipelines for the detection of small insertion/deletions and SNV.

**Table 2:**
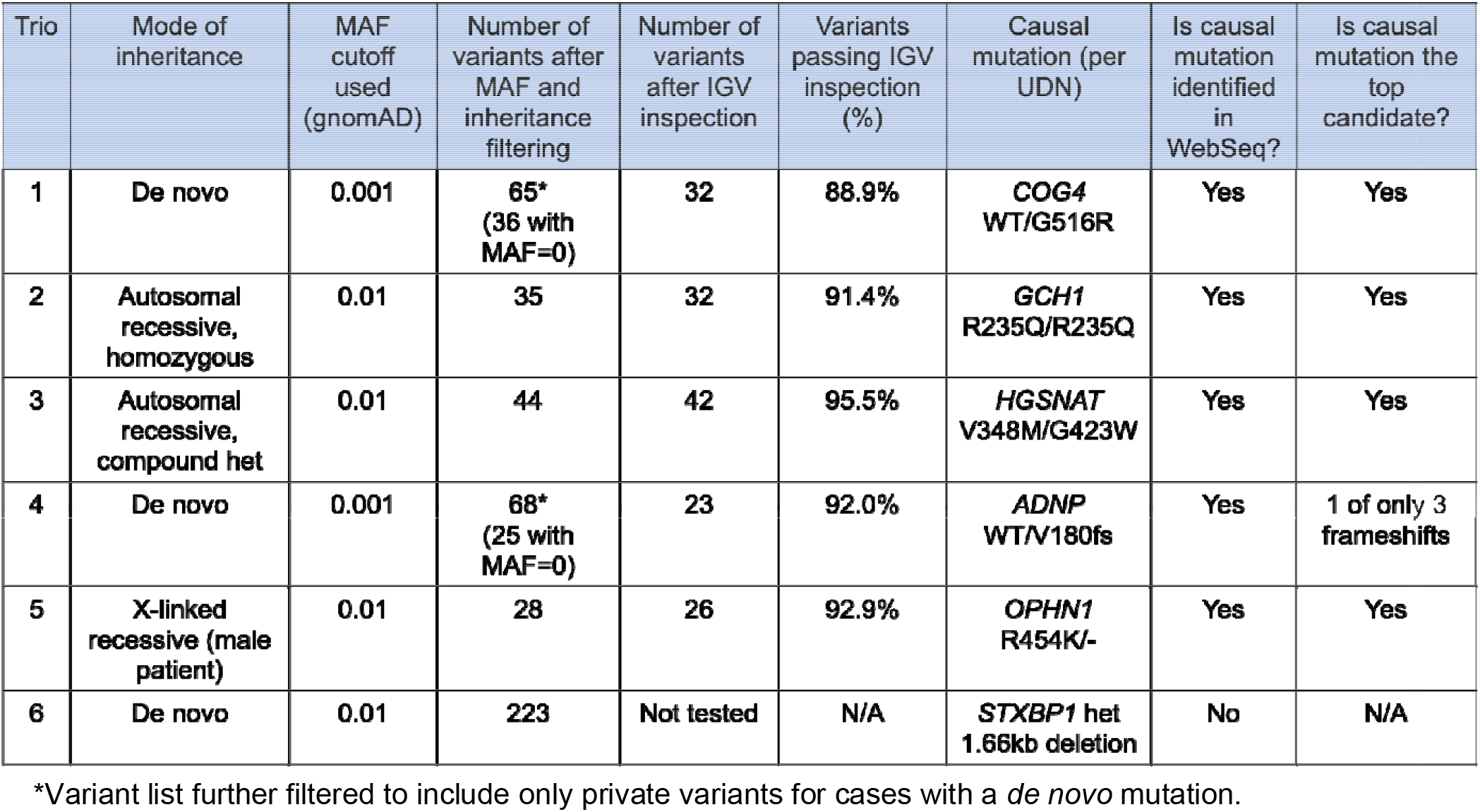
Validating WebSeq functionality using previously analyzed data from UDN (Undiagnosed Disease Network)

| Trio | Mode of inheritance | MAF cutoff used (gnomAD) | Number of variants after MAF and inheritance filtering | Number of variants after IGV inspection | Variants passing IGV inspection (%) | Causal mutation (per UDN) | Is causal mutation identified in WebSeq? | Is causal mutation the top candidate? |
| --- | --- | --- | --- | --- | --- | --- | --- | --- |
| 1 | De novo | 0.001 | 65*<br>(36 with MAF=0) | 32 | 88.9% | COG4<br>WT/G516R | Yes | Yes |
| 2 | Autosomal recessive, homozygous | 0.01 | 35 | 32 | 91.4% | GCH1<br>R235Q/R235Q | Yes | Yes |
| 3 | Autosomal recessive, compound het | 0.01 | 44 | 42 | 95.5% | HGSNAT<br>V348M/G423W | Yes | Yes |
| 4 | De novo | 0.001 | 68*<br>(25 with MAF=0) | 23 | 92.0% | ADNP<br>WT/V180fs | Yes | 1 of only 3 frameshifts |
| 5 | X-linked recessive (male patient) | 0.01 | 28 | 26 | 92.9% | OPHN1<br>R454K/- | Yes | Yes |
| 6 | De novo | 0.01 | 223 | Not tested | N/A | STXBP1 het<br>1.66kb deletion | No | N/A |

\*Variant list further filtered to include only private variants for cases with a *de novo* mutation.

## Discussion

Given the increasing application of human genomics data to understanding human health, we cannot afford to restrict the number of researchers who can conduct genomic data analyses. Heavy reliance on command line to build popular bioinformatics toolkits such as ANNOVAR^1^, KING^9^, PLINK^5,6^ keeps users without command line skills from appreciating and using these extremely valuable tools. Genomic data analytics tools need to be user-friendly, modern, easy to use, portable, and secure. This is the motivation behind developing WebSeq, a one-stop user-friendly web interface for genomic data analytics.

Building web interfaces for command line toolkits is not new in the bioinformatics community. For example, the creators of the popular variant annotation toolkit ANNOVAR^1^ also built a web-based version-wANNOVAR^3^. wANNOVAR’s^3^ modern web-based interface is very easy to use and performs some of the more popular functionalities of ANNOVAR^1^. However, it enforces a file size restriction on VCF files that is incompatible with current WES and WGS methodologies. This is in part due to the conversion process being performed on wANNOVAR^3^ server. We learn from this effort and conduct all variant annotation in WebSeq on the user’s computer locally, solving the file size restriction problem. Since users no longer need to send their data to remote servers for analysis, conducting local analyses also keeps patient data safe and secure.

A similar effort was made by the developers of PLINK^5,6^ who recently built gPLINK^35^. gPLINK^35^ is a Java based Desktop interface that provides a simple interface to the more commonly used PLINK^5,6^ commands^35^. While users must utilize command line for installation and debugging purposes, gPLINK^35^ works on all major operating systems. However, WebSeq utilizes a variety of popular bioinformatics toolkits that do not enjoy such cross-platform support. To ensure that all modules within WebSeq’s modern web-based framework were also capable of running on all operating systems, we utilized Docker^4^. Docker^4^ offers operating-system level virtualization to deliver software in bundled packages called containers. We ship Docker^4^-enabled WebSeq containers with all necessary programming languages, software, libraries, configuration files preinstalled, so researchers can start their analyses with ease.

WebSeq allows researchers to use sophisticated genomics analytics pipelines without experience with Bash or command line tools. WebSeq is also hosted locally which allows for secure analyses without sharing protected data with servers like wANNOVAR^3^ or GATK^7^ Cloud. WebSeq is made even more powerful by adding cross-platform support using Docker^4^ that enables ease of installation and use. We hope that WebSeq will contribute to the growing suite of accessible and streamlined web-based bioinformatics tools and support intuitive analysis of human genomics data for its users.

## Software Details

Project name: WebSeq

License: free for personal, academic, and non-profit use

Application available at: http://bitly.ws/g6cn

Application Sizes:

- WebSeq Lite: 5GB
- WebSeq: 50GB
- WebSeq Pro: 100GB

## Acknowledgements

The authors wish to acknowledge helpful input during the development and testing stages of this toolkit, provided by Dr. Marcela Moncada-Velez (Rockefeller University), Drs. Ruben Martinez-Barricarte, Tyne Miller-Fleming, Rui Chen, and Qiang Wei (Vanderbilt University Medical Center), Zhiming Mao and Julia Ryan (Johns Hopkins University). We would also like to thank Dr. Kai Wang (University of Pennsylvania) for allowing us to integrate ANNOVAR into this toolkit for non-profit, academic use. The authors received funding support for this work from the Provost’s Undergraduate Research Award at Johns Hopkins University, The Johns Hopkins Institute for Data-Intensive Engineering and Science, and the Vanderbilt Human Immune Discovery Initiative (HIDI). We thank all of the individuals and their families for their participation in this study. Research reported in this paper was supported by the NIH Common Fund through the Vanderbilt University Medical Center (U01HG007674) (Vanderbilt University Medical Center). The content is solely the responsibility of the authors and does not necessarily represent the official views of the National Institutes of Health.

